# Normal aging increases white matter microglial reaction and perivascular macrophages in the microcebe primate

**DOI:** 10.1101/2025.03.26.645519

**Authors:** Léo Dupuis, Lolie Garcia, Fanny Petit, Suzanne Lam, Helène Hirbec, Jean-Luc Picq, Delphine Boche, Marc Dhenain

## Abstract

As populations age, the incidence of neurodegenerative disorders is rising. Early age-related neuropathological changes are the breeding ground for the development of these disorders. Microglia are the resident macrophages of the central nervous system. They play crucial roles in maintaining brain homeostasis, yet their age-related changes remain not fully understood. While age-related microglia changes have been strongly characterized in rodents, studies in non-human primates are scarce. The microcebe primate (mouse lemur (*Microcebus murinus*)) is widely used as a model for cerebral aging and to investigate age-related neurodegenerative processes. HLA-DR is a major histocompatibility class II cell surface receptor which presents antigens to cells of the immune response. It is a major marker of microglia reaction. In this study, we explored microglia in the whole brains of middle-aged and old microcebes using HLA-DR immunolabeling. We analyzed microglial morphology and quantified HLA-DR+ cell density and protein expression. A wide range of microglial morphologies was observed in the white matter, including thin processes microglia, “rod-like” elongated and polarized shape, hypertrophic, and amoeboid microglia. Aging was associated with increased HLA-DR+ microglial expression in the white matter while very few HLA-DR+ microglia were observed in the parenchyma of cortical gray matter regions. A second finding was the higher number of HLA-DR+ perivascular macrophages in old animals. This study in a primate outlines that, in the absence of neurodegenerative processes, the most prominent signs of age-related microglia/macrophage changes are region-specific and concern white matter and perivascular regions. This emphasizes the need to target these regions to prevent cerebral aging.

**Main Points:**

- HLA-DR+ microglia in *Microcebus murinus* primate show diverse morphologies and increase with age in white matter.
- HLA-DR+ perivascular macrophage load increase in aged animals.

**Table of Contents Image (TOCI):** 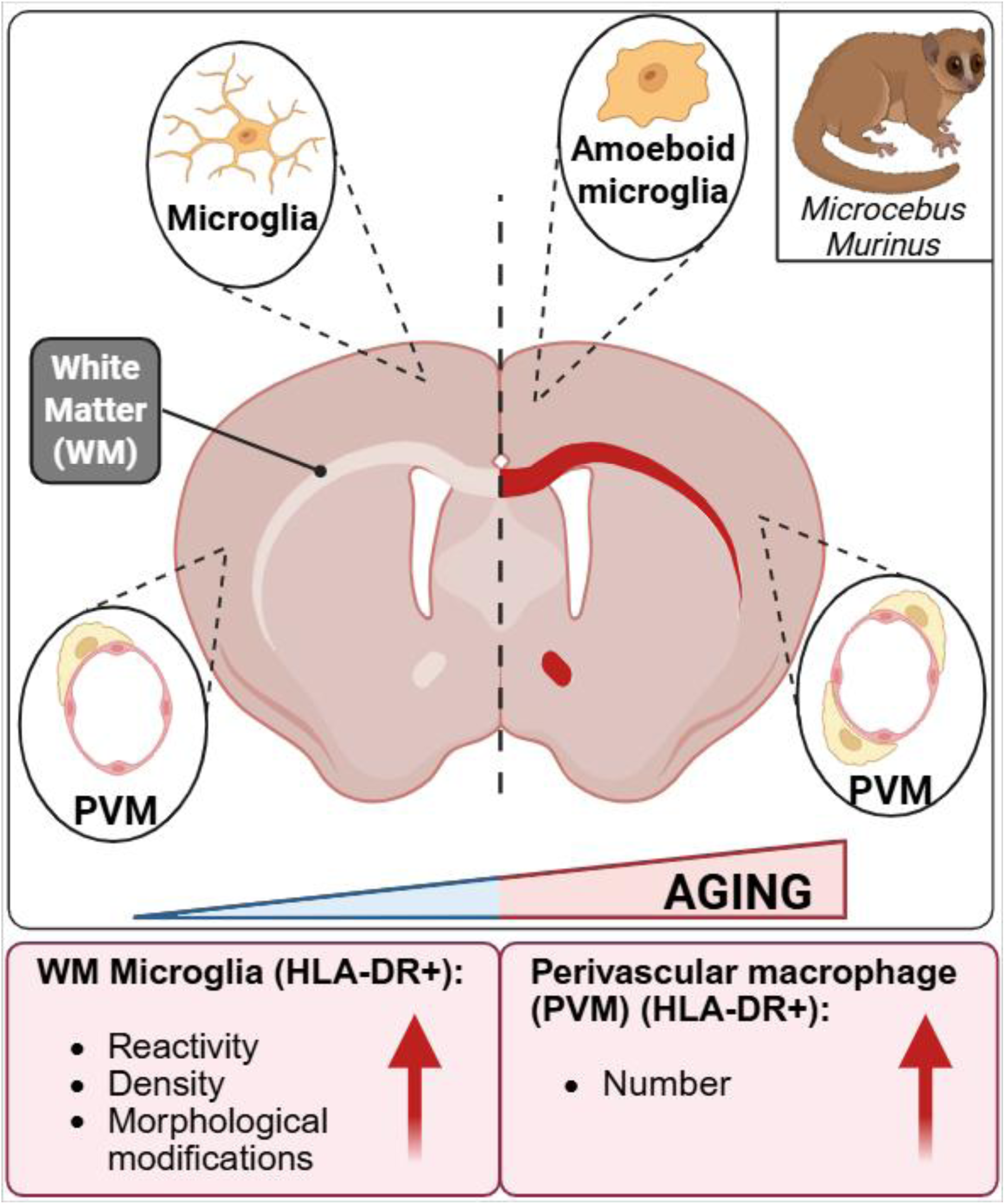

## 1. Introduction

Over the past century, human life expectancy has risen substantially, accompanied by a higher likelihood of brain changes that can result in neurodegenerative disorders as well as mild cognitive and motor impairments affecting daily living. Examination of brain tissue from elderly individuals frequently reveals pathological changes, even in those without clinical symptoms. These changes are the breeding ground for the development of neurodegenerative processes.

Microglia are the resident macrophages of the central nervous system (CNS), comprising approximately 10% of brain cells (Paolicelli et al., 2022). They play crucial roles in maintaining brain homeostasis. Several studies have outlined that during aging, microglial density, morphology or activation status change (Edler et al., 2021). In humans microglia labeling is reported in white matter of adult subjects (Mattiace et al., 1990) and an age-related increased reactivity is shown in this region (Gefen et al., 2019; Mattiace et al., 1990; McGeer et al., 1988; Rogers et al., 1988; Sobel & Ames, 1988); while in cortical regions, aging does not seem to affect microglia (Mattiace et al., 1990; Rogers et al., 1988; Sobel & Ames, 1988), unless amyloid plaques, one of the lesions of Alzheimer’s disease occur (Boche & Nicoll, 2020). Observational studies in humans are hampered by difficulties due to comorbidities, agonal status, and post-mortem delays. Also, experimental studies can hardly be performed in humans. Many studies investigating microglial changes during aging have thus been conducted in rodents. However, the relevance of rodent studies to human microglia has been challenged, due to differences in life expectancy, microglial dynamics and functional roles in physiological conditions (Boche & Gordon, 2021). Nonhuman primates share greater genetic and physiological similarities with humans compared to rodents. They thus offer a valuable perspective on microglial status during aging.

The microcebe (mouse lemur (*Microcebus murinus*)) is among the smallest nonhuman primates. Measuring approximately 12 cm in length, it has a relatively short lifespan for a primate of around 12 years. Some old animals can exhibit age-related cognitive decline (Picq et al., 2012) as well as amyloid and tau lesions (Kraska et al., 2011). This model is recognized to study aging and neurodegenerative diseases (Lam et al., 2021; Languille et al., 2014), particularly for exploring how cerebral aging is influenced by biological factors and conditions such as chronic caloric restriction (Pifferi et al., 2018) or diabetes (Djelti et al., 2016).

Given the importance of the microcebes for biomedical research, it is critical to increase our knowledge on microglia in this species. In this study, we evaluated age-related changes of microglia in microcebes, exploring the expression of HLA-DR protein. HLA-DR is a major histocompatibility class (MHC) II cell surface receptor that presents antigens to cells of the immune response. It is involved in the non-self-recognition and upregulated in inflammation (Sloane et al., 1999; Styren et al., 1990). HLA-DR was also identified as a genetic risk factor for Alzheimer’s disease (Jones et al., 2010) and its expression in humans is associated with the presence of dementia and Alzheimer’s pathology (Minett et al., 2016). Here, we described microglial morphology and macrophage populations (meningeal and perivascular), as well as age-related microglial regional changes in the microcebe.

## 2. Materials and methods

### 2.1. Microcebes

A total of seventeen micocebes were studied with an age range of 1.0 to 11.5 year-old (mean±SD: 8.8 ± 1.1 years) comprising four animals in middle-age (n=3 males, n=1 female, 3.1 ± 2.0 year-old), and 13 aged animals (n=6 males, 7 females, 10.4 ± 1.5 year-old; Supplementary table 1). The animals were bred in the animal facility of the Molecular Imaging Research Center, CEA, Fontenay-aux-Roses. Microcebes were kept in an enriched environment, temperatures between 24-26°C, relative humidity of 55%, and seasonal illumination (summer: 14h of light, 10h of darkness; winter: 10h of light, 14h of darkness). Food consisted of fresh apples and a handmade blend of bananas, cereals, eggs, and milk. Water supply for animals was freely accessible. None of the animals had ever taken part in invasive research or pharmaceutical trials before. The 13 old animals and one middle-aged animal were sacrificed specifically for the study without presenting any identified pathology expected to lead to imminent spontaneous death. One middle-aged animal was euthanized following tail infection to shorten animal suffering. The brains of two middle-aged animals were sampled shortly after their spontaneous death from accident and sudden death. Magnetic resonance imaging were performed on all animals to rule out obvious cerebral pathology. MRI scans did not reveal any vascular pathology or severe atrophy process. Post-mortem autopsy was performed on each animal to evaluate hidden pathologies. Finally, histological staining for amyloid-β (Aβ) and Tau was performed on all the brains to rule out Alzheimer like pathologies. No peripheral chronic diseases were detected, and only one animal exhibited amyloidosis; this individual was excluded from the cohort and analyzed as a case report.

### 2.2. Ethics

All experimental procedures were performed in compliance with the European Union directive on the protection of animals used for scientific purposes (2010/63/EU) and French regulations (Rural Code R214/87-131). They were approved by a local ethics committee (CETEA-CEA DSV IdF) as well as by the French Ministry of Education and Research (authorization A14_035 and A17_083). All efforts were made to minimize animal suffering and animal care was supervised by veterinarians and animal technicians skilled in microcebe healthcare and housing.

### 2.3. Perfusion and tissue preparation

The sacrificed animals were killed with an intraperitoneal injection of pentobarbital sodium (0.1ml/100g; Exagon, Axience). Twenty minutes before any incision, a subcutaneous administration of buprenorphine (0.1mg/100g; Vétergésic®) was performed for analgesia. Animals were perfused intracardiacally with 0.1M PBS. The brain was post-fixed in 4% paraformaldehyde for 48h at 4°C., transferred in a 15% sucrose solution for 24h and in a 30% sucrose solution for 48h at 4°C for cryoprotection. Serial coronal sections of 40µm were performed with a microtome (SM2400, Leica Microsystem) and stored at -20°C in a storing solution (glycerol 30%, ethylene glycol 30%, distilled water 30%, phosphate buffer 10%). Free-floating sections were rinsed in a 0.1M PBS solution (10% Sigma-Aldrich® phosphate buffer, 0.9% Sigma-Aldrich® NaCl, distilled water) before use.

### 2.4. Immunohistochemistry

A key resource table is provided in Supplementary Table 2. Microcebe brains were stained for microglia using the Human Leukocyte Antigen – DR isotype (HLA-DR, clone TAL.1B5, Dako M0746) and for Alzheimer’s hallmarks with amyloid-β (Aβ) (clone 4G8, Biolegend 800706) and tau (clone AT8, Thermo MN1020B) antibodies.

For HLA-DR staining, a pretreatment was applied with EDTA 1X citrate (Diagnostic BioSystems®) at 95°C followed by 0.5% Triton X-100/0.05M Tris-HCl Buffered Saline solution (TBS). Tissues were then incubated in BLOXALL (Impress kit, Eurobio Scientific®) to inhibit endogenous peroxidases. Sections were then incubated at 4°C in a 3% BSA/TBST solution with the HLA-DR antibody (dilution 1/250) for 48h followed by the Impress Kit (Impress kit, Eurobio Scientific®). For 4G8 labeling, a pretreatment with 70% formic acid (VWR®) was performed. For both 4G8 and AT8 staining, tissues were incubated at 4°C with 4G8 (dilution 1/350) or AT8 (dilution 1/500) antibody in a 3%BSA/PBST (PBS-Triton 0.5%) solution. Incubation with the appropriate biotinylated secondary antibody followed by the avidin-biotin complex solution (ABC Vectastain kit, Vector Laboratories®) was then applied.

Revelation was performed using the DAB Peroxidase Substrate Kit (DAB SK4100 kit, Vector Laboratories®). Sections were mounted on Superfrost Plus slides (Thermo-Scientific®). For the 4G8 and AT8 labeling, a cresyl violet counterstain was performed. All sections were dehydrated and the slides mounted with the Eukitt® mounting medium (Sigma Aldrich).

### 2.5. Image acquisition and quantification

Images were acquired at 20× using an Axio Scan.Z1 (Zeiss®). Each section was extracted individually in the .czi format using the Zen 2.0 (Zeiss®) software. Images were then imported in QuPath v0.4.3software (the University of Edinburgh, UK (Bankhead et al., 2017)). Quantification of the staining relied on a threshold manually established, validated across multiple images, and used similarly across all animals. An average of four sections per animal, selected at different anatomical levels, were analyzed to provide a comprehensive overview of the whole brain. Regions of interest (ROIs) were manually segmented based on a microcebe atlas (Nadkarni et al., 2019). Six ROIs were manually outlined from each brain to include: the corpus callosum, optic chiasma, hippocampus, cingulate cortex, parietal cortex, and entorhinal cortex. The analysis was performed blind to the age of the animal. One animal was excluded from the quantification of the hippocampus, entorhinal cortex, and parietal cortex due to tissue damage.

Cell counting was performed using the “Analyze - Cell Detection – Fast Cell Counts” function in QuPath. Results show average values from different sections for each animal. Perivascular macrophages were manually counted within each ROI. Each HLA-DR+ cells inside or near blood vessels was considered as perivascular macrophages. Blood vessels in each region were manually counted using Cresyl Violet staining. The density of blood vessels in each area was calculated as explained above. The number of perivascular macrophages or blood vessels was calculated using the formula - (cell or blood vessel count / area)×1,000,000 - yielding the number of macrophages / blood vessels per mm².

### 2.6. Luxol Fast Blue Staining

Myelin fibers were detected using Luxol Fast Blue staining (Fisher Scientific®). Mounted sections (40µm) were placed in 95% ethanol and then incubated in 0.1% Luxol fast blue. Then, slides were washed and placed in 0.05% lithium carbonate before final wash. Slides were desiccated in 100% ethanol and washed twice in xylene. All slides were cover-slipped using Eukitt® (Sigma Aldrich). Images were acquired at 20x using 3Dhistch Panoramic P250 Flash II scanner.

### 2.7. Statistical analysis

All statistical analyses were conducted using RStudio (RStudio Team, 2020) and R version 4.4.2. Differences between groups within different regions were assessed using non-parametric ANOVA based on rank value (nparLD package (Noguchi et al., 2012)). Then, homogeneity of variances was assessed using Levene’s test (Levene, 1960) (car package (Fox & Weisberg, 2011)). If the assumption of equal variances was met, pairwise permutation t-tests (pairwise.perm.t.test function, 1000 permutations with Benjamini-Hochberg correction) were applied (RVAideMemoire package (Hervé, 2020)). If the assumption of equal variances was violated, a rank-based non-parametric t-test was applied (npar.t.test function from the nparcomp package (Derrick et al., 2020)). For all statistical tests, p-values were adjusted for multiple comparisons using the Benjamini-Hochberg method (Benjamini & Hochberg, 1995). The difference in HLA-DR surface area between anterior and posterior sections was assessed using the non-parametric Wilcoxon signed-rank test (Rcmdr package (Fox & Weisberg, 2011) in R). A significance threshold of p < 0.05 was applied for all tests.

## 3. Results

### 3.1. Diversity of HLA-DR immunoreactivity in microcebe brains

Macroscopic observation of HLA-DR immunoreactivity revealed strong labeling in some white matter regions such as the corpus callosum or optic chiasma and a sparse labeling in gray matter (Fig. 1). White matter staining was highly heterogeneous amongst the animals, with low or no staining in middle-aged animals (Fig. 1A) and a range from no to high staining in old animals (Fig. 1B-D).

**Figure 1.**
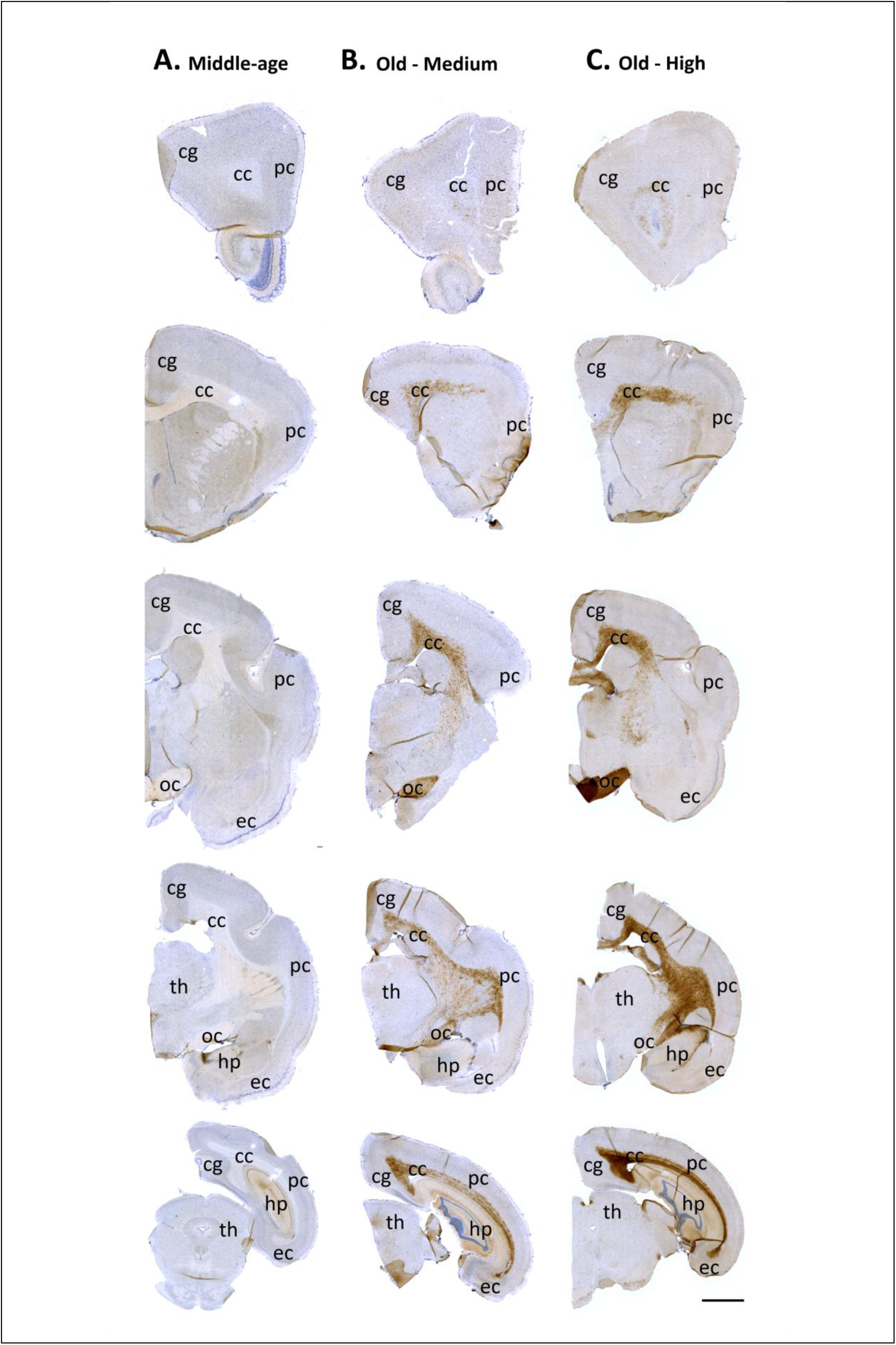
HLA-DR immunoreactivity throughout the brain of middle-aged and old microcebes. HLA-DR immunoreactivity was observed mainly in white matter areas (corpus callosum and optic chiasma), while the staining was low to absent in gray matter. Middle-age brain displaying no HLA-DR immunoreactivity in both white and gray matter (A). Old microcebes displaying a medium (B) or strong (C) HLA-DR immunoreactivity mainly in white matter areas (B). Scale bars = 2 mm. cg for cingulate cortex, pc for parietal cortex, ec for entorhinal cortex, cc for corpus callosum, oc for optic chiasma, th for thalamus.

Three of the four middle-aged microcebes and four out of the thirteen old animals showed no HLA-DR+ expression (Fig. 2A). One middle-aged microcebe and four older animals displayed a low HLA-DR load (Fig. 2B) characterized by microglia with thin processes and low HLA-DR expression (Fig. 3A-C). Most of them displayed a “rod” shape characterized by an elongated soma with polarized processes mainly in the apical and basal ends of the cell aligned with myelin fibers (Fig 3D-F). Three old animals exhibited a medium HLA-DR expression (Fig. 2C), marked by microglia with thicker processes and enlarged cell body (Fig. 3H-I). Some microglia exhibited bulbous ending processes (Fig. 3G). Finally, two old microcebes demonstrated a high HLA-DR expression (Fig. 2D), predominantly characterized by intensely labeled microglia with amoeboid shape (big round soma) with or without thin processes (Fig. 3K-L), indicative of heightened reactivity. More rarely, some hypertrophic microglia with a highly ramified “bushy” shape were also detected in medium and high HLA-DR immunoreactivity profiles (Fig. 3M).

**Figure 2.**
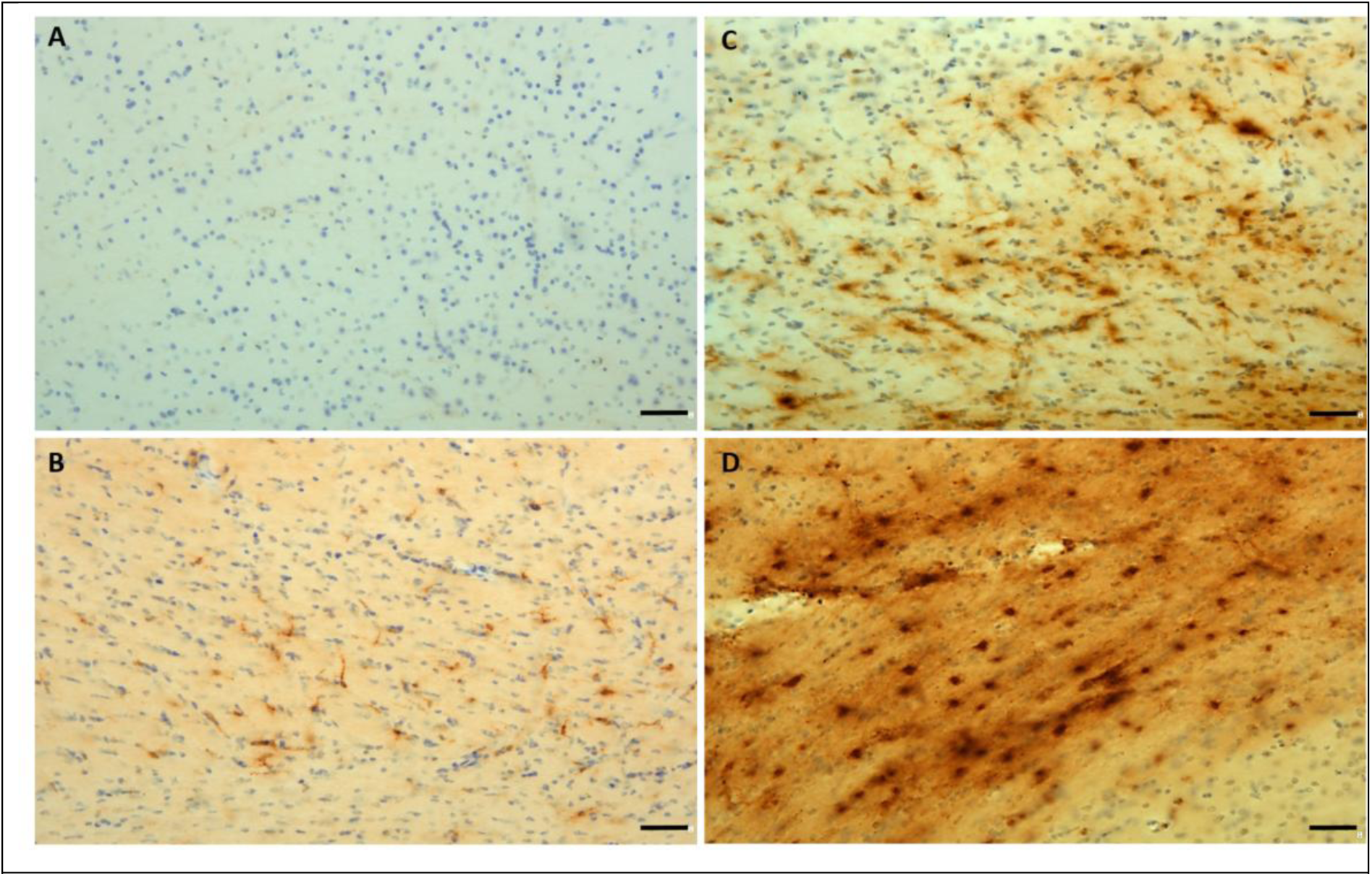
HLA-DR immunoreactivity throughout the corpus callosum of microcebes during aging. Absence of HLA-DR immunoreactivity (A). Low (B); medium (C); high (D) HLA-DR immunoreactivity. Scale bar = 40µm.

**Figure 3.**
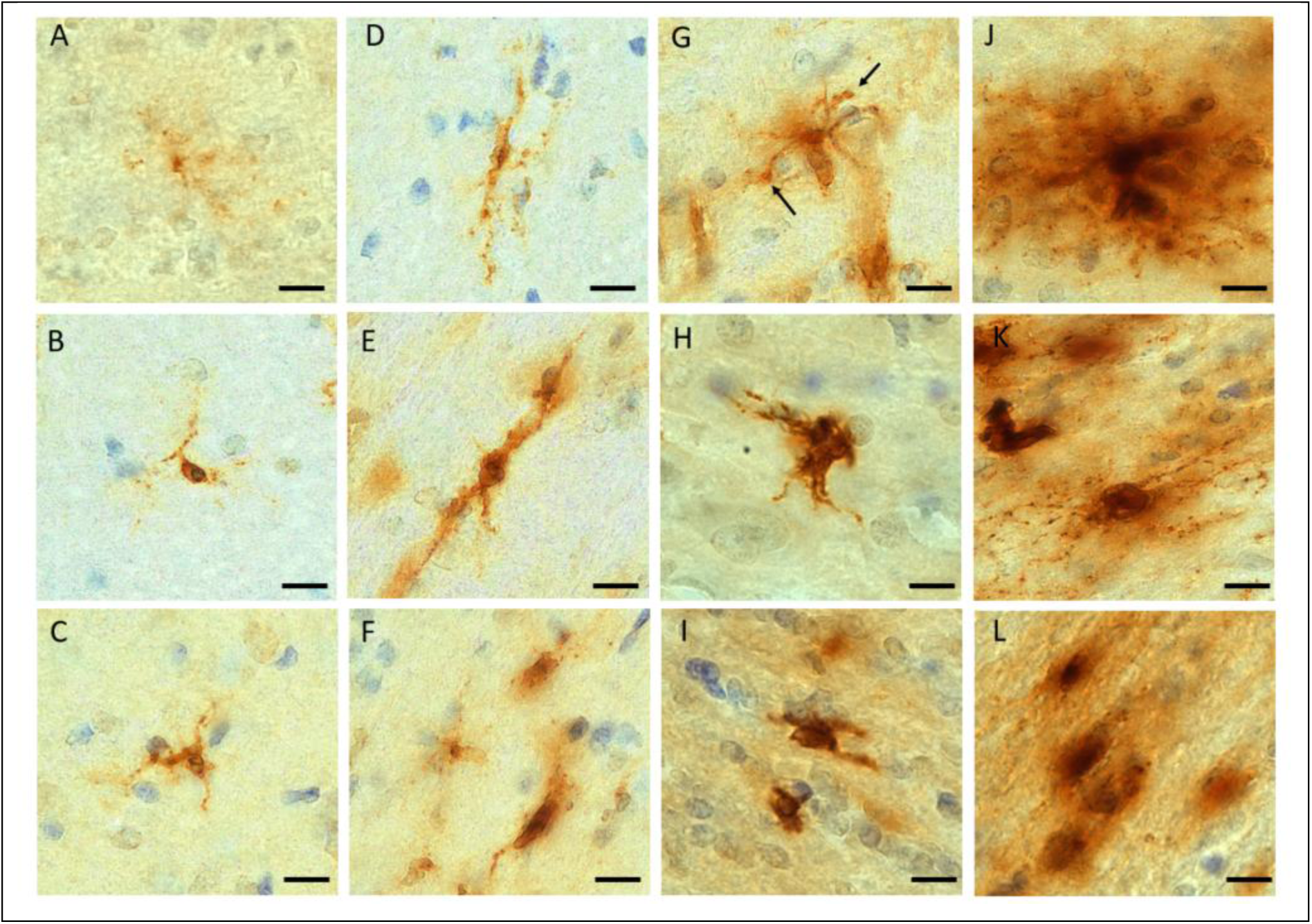
Diversity of HLA-DR+ microglia in the white matter. Microglia with thin processes (A-C). Rod shaped microglia with a long cellular cell body from which emerged bipolar processes along the myelin fibers (D-F). Microglia with bulbous ending processes (black arrow, (G)). Microglia with an enlarged cell body as well as short and thick cellular processes (H-I). Hypertrophic microglia highly ramified (J). Amoeboid microglia with enlarged cell body and thin or absent processes (K-L). Scale bar = 10µm.

In the cortical gray matter, HLA-DR immunoreactivity was less prominent than in white matter. HLA-DR also labeled the perivascular macrophages within the blood vessel walls (Fig. 4A-B). Most HLA-DR+ cells in the parenchyma were located in white matter (Fig. 4C), as highlighted by the Luxol blue stain (Fig. 5B). Very few HLA-DR+ microglia were observed in other cortical gray matter regions (Fig. 4D). Additionally, macrophages were detected in the meninges (Fig. 4E) and the choroid plexus (Fig. 4F).

**Figure 4.**
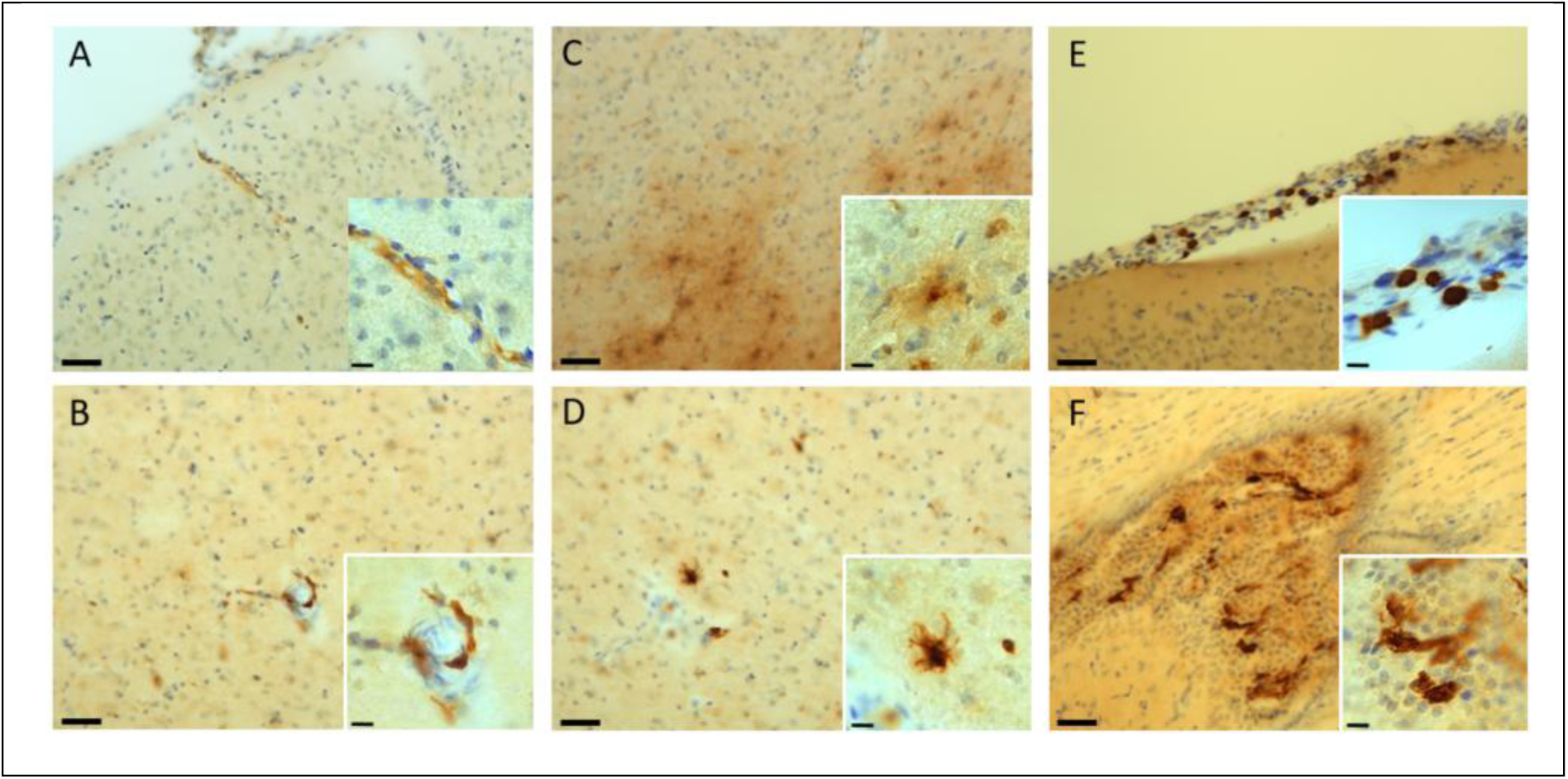
HLA-DR+ cells in cortical gray matter, meninges and choroid plexus. Perivascular macrophages (A-B). Microglia in the deep layers of the cortex near the white matter (C). Immune cells in the upper layers of the cortex (D). Meningeal macrophages (E). Macrophages in the choroid plexus (F). Scale bar 40µm and 10µm in the zoomed box.

**Figure 5.**
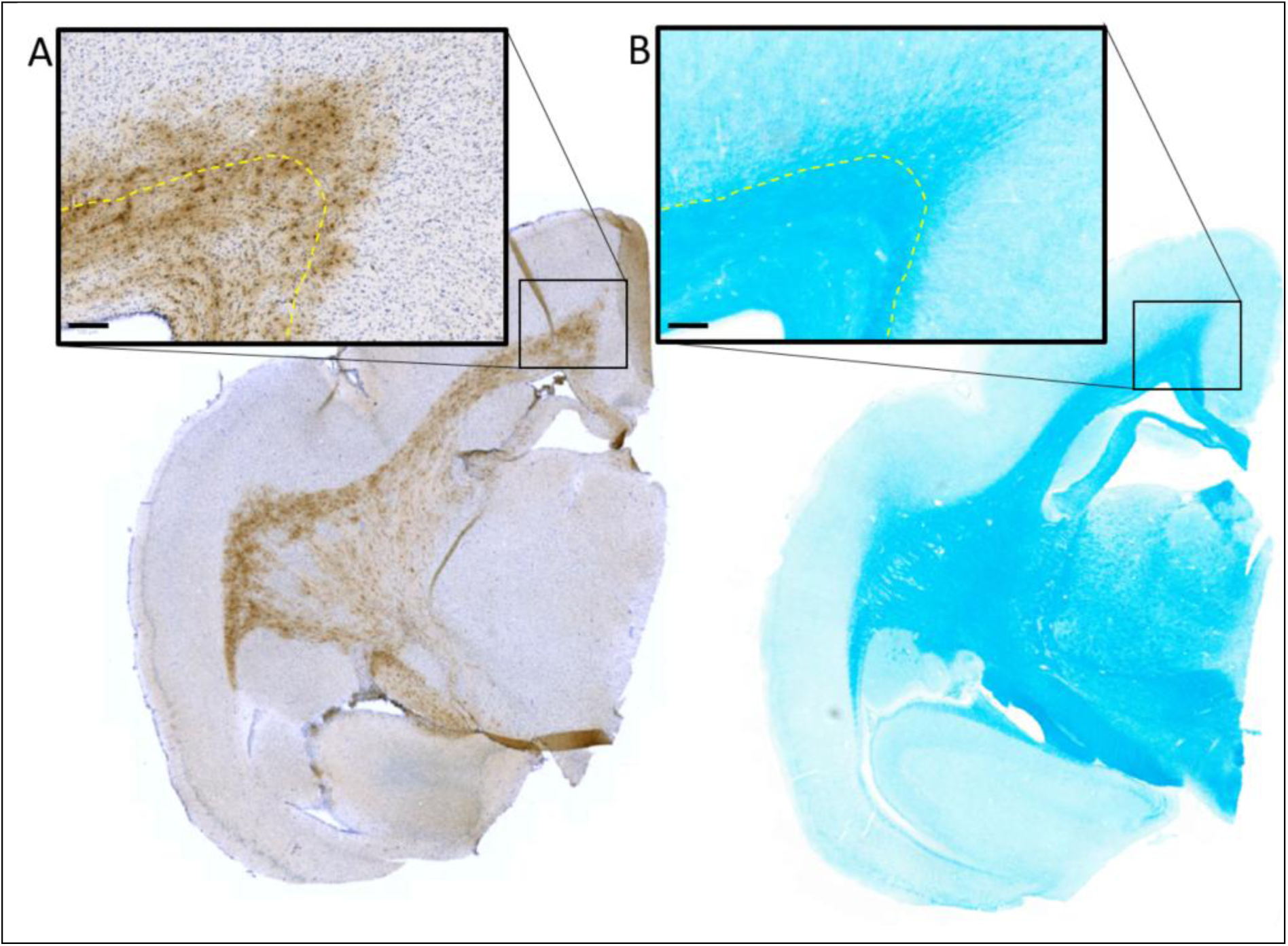
HLA-DR+ microglia in white matter. (A) HLA-DR staining in the corpus callosum (under the yellow line) and in the deep layer of the cortex (above the yellow line). (B) Luxol Fast Blue stain to visualize the myelin fibers in the corpus callosum (under the yellow line) and in the deep layer of the cortex (above the yellow line). Scale bar = 100µm.

### 3.2. Quantitative analysis of age-related HLA-DR labeling

To further investigate age-related changes of HLA-DR expression, quantitative analysis of HLA-DR+ protein load or cells was performed. Analysis of HLA-DR protein load revealed a significant higher expression in old animals in the corpus callosum and optic chiasma compared to middle-aged (Fig. 6A, corpus callosum: p = 0.006; optic chiasma: p = 0.012). HLA-DR+ cell showed a significant increase of cell density in the corpus callosum but not in the optic chiasma in old microcebes compared to middle-aged ones (Fig. 6B, corpus callosum: p = 0.022; optic chiasma: p = 0.22). As illustrated in Fig. 1C, the age-related HLA-DR protein load in the corpus callosum was significantly higher in posterior compared to anterior brain sections (Fig. 6C, p = 0.00049).

**Figure 6.**
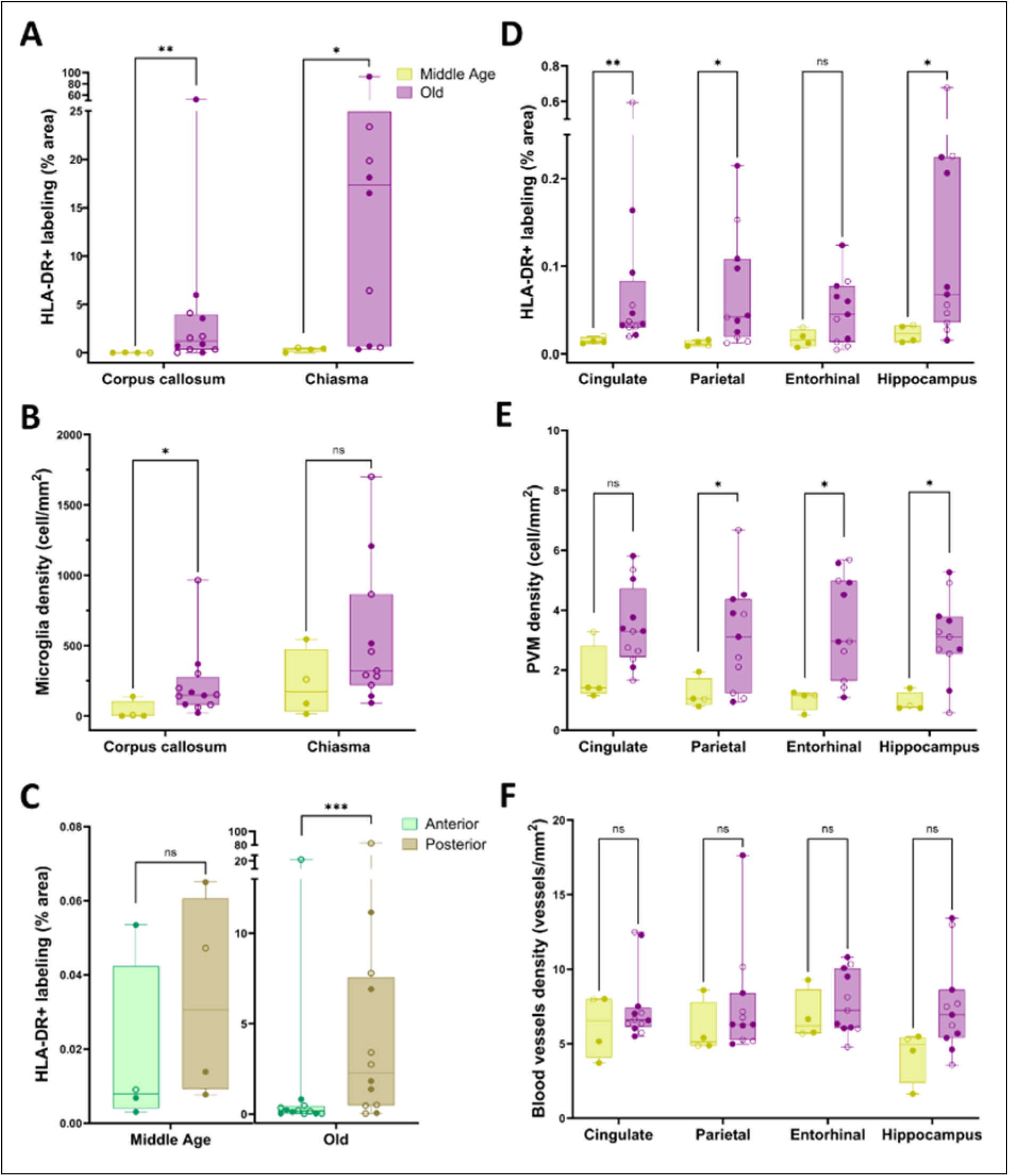
Quantification of HLA-DR immunoreactivity in the white and gray matter of middle-aged and old microcebes. (A) HLA-DR protein load in the white matter revealed an age-related increase of microglial immunoreactivity in the corpus callosum and the chiasma optic (p<0.001 - non-parametric ANOVA group comparison; p<0.05 with pairwise permutation t-tests and Benjamini-Hochberg (BH) p-values adjustment). (B) HLA-DR+ cells density (cells/mm^2^) in the white matter showed an age-related increase in the corpus callosum but not in the optic chiasma (p<0.001 - non-parametric ANOVA group comparison; p<0.05 with pairwise permutation t-tests and BH p-values adjustment). (C) HLA-DR protein load in the corpus callosum was higher in the posterior brain section of old microcebes (p<0.05; Wilcoxon signed rank test) but not in middle age animals (p>0.05). (D) HLA-DR protein load revealed an age-related increase in microglia immunoreactivity in the cingulate cortex, parietal cortex and hippocampus (p<0.001 - non-parametric ANOVA group comparison; p<0.05 with pairwise permutation t-tests and BH p-values adjustment) but not in the entorhinal cortex (p>0.05). (E) Density of perivascular macrophages (cells/mm^2^) increased with aging in the parietal and entorhinal cortex as well as in the hippocampus (p<0.001 - non-parametric ANOVA group comparison; p<0.05 with rank-based non-parametric t-test and BH adjustment) but not in the cingulate cortex (p>0.05). (F) Blood vessels density (vessel/mm^2^) was similar between middle-aged and old microcebes (p>0.05 with pairwise permutation t-tests and BH p-values adjustment). Microcebe’s sex is represented with a full circle for males and an empty circle for females. *p < 0.05; **p < 0.01; ***p < 0.001; ns= non-significant.

The HLA-DR protein load was higher in white matter compared to cortical regions. Analysis in the cortical areas revealed an increased expression in the cingulate and parietal cortex but not in the entorhinal cortex of old compared to middle-aged animals (Fig. 6D, cingulate: p = 0.002; parietal: p = 0.016; entorhinal: p = 0.092; hippocampus: p = 0.032). HLA-DR expression was also higher in the hippocampus of old animals (Fig. 6D, p = 0.032). Perivascular macrophages counting uncovered a significant increase of cell density in old compared to middle-aged animals in parietal and entorhinal cortex as well as in the hippocampus (Fig 6E, parietal p = 0.037, entorhinal cortex p = 0.037, hippocampus: p = 0.037). The vascular density was not modified in these regions (Fig. 6F, Cingulate: p = 0.74, parietal p = 0.53, entorhinal cortex p = 0.71, hippocampus: p = 0.37). These results indicate that the majority of HLA-DR immunoreactivity observed in the gray matter of aged animals was primarily due to an increase in perivascular macrophages. Notably, in the cingulate cortex, no change in the number of perivascular macrophages was detected, despite a pronounced age-related rise in HLA-DR protein load. In this region the observed increase in reactivity was mainly attributed to myelin-associated microglia located at the border of the corpus callosum as seen on Luxol staining (Fig. 4B). Of note, sex differences were analyzed in older animals revealing no significant differences between males and females (not shown).

### 3.3. Lack of microglia reactivity close to amyloid deposits in an old microcebe

One aged animal that exhibited diffuse amyloid plaques in the cortical region (Fig 7A), was assessed to investigate whether amyloid deposits could alter microglial reactivity. As illustrated in Fig. 7, despite the presence of amyloid plaques (Fig. 7A), no HLA-DR microglia were detected (Fig. 7B).

**Figure 7.**
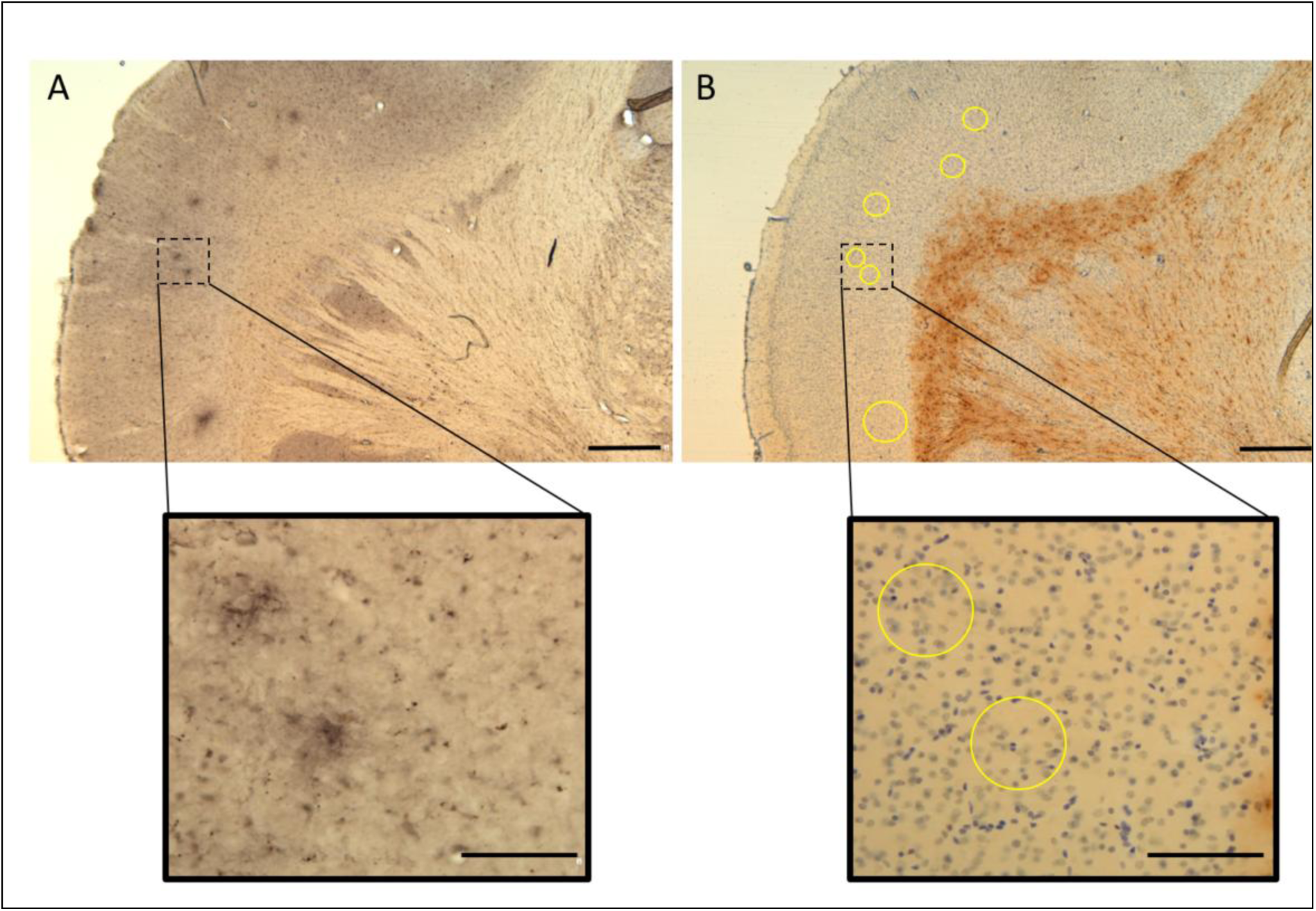
Cortical HLA-DR+ microglia were not detected despite the presence of amyloid plaques. (A) 4G8 labeling showing diffuse amyloid plaques in the parietal cortex. (B) HLA-DR labeling showing the absence of microglia near the amyloid plaque. Yellow circle = projection of plaque localization. Scale bar 500µm and 100µm in the zoomed box.

## 4. Discussion

### 4.1. Diversity of HLA-DR microglial immunoreactivity in microcebe brains

Previous studies evaluating microglia in microcebe brains have reported microglia, evidenced by anti-ferritine antibody, close to amyloid plaques in the cortex with their presence being decreased by anti-amyloid vaccines (Trouche et al., 2023). However, despite extensive efforts, no study succeeded in labeling homeostatic microglia in the brain of microcebes. Also, no study characterized microglia in the aging brain. Some studies have focus on microglia in the spinal cord. They showed reactive and amoeboid microglia in spinal cord injured regions while those in intact regions remained ramified (Poulen et al., 2021). Here we decided to investigate for the first time HLA-DR expression in the microcebe brain across aging.

We reported that HLA-DR expression is prevalent in the white matter, with few cells detected in the gray matter. Age-related change of HLA-DR microglial expression in the white matter was associated with a pronounced heterogeneity in the number of HLA-DR+ microglia among the old animals. Also, the morphology status of predominant microglia was associated with the immunoreactivity levels: Animals with low HLA-DR expression profile had ramified or “rod” shape microglia, the latter morphology having been reported in the white matter without any pathological context (Holloway et al., 2019). Those with medium HLA-DR expression had microglia with thicker processes and enlarged cell body, while the animals with high HLA-DR expression presented amoeboid morphology of microglia. In addition, “bushy” microglia were detected in animals with medium and high expression.

Previous studies have described age-related white matter atrophy in microcebes (Picq et al., 2012; Sawiak et al., 2014) suggesting that white matter is a region particularly vulnerable to aging effects in this species. Mouse studies have highlighted that debris released by aging myelin sheaths are cleared by microglia (Conde & Streit, 2006; Safaiyan et al., 2016), and we can hypothesize that a similar event happens in microcebes driving increased microglial activation in the white matter with aging. In addition to aging processes, we cannot rule out that other pathological states could lead to microglia activation in some animals, explaining some of the reported microglial heterogeneity observed in old microcebes, with some individuals showing high HLA-DR+ expression of microglia in the white matter, while others exhibited little to no expression of HLA-DR. Indeed, aging in microcebes is also associated with age-related pathologies, including sarcopenia (Hamalainen et al., 2015) or diabetes (Djelti et al., 2016). The functional impact of microglial activation remains to be evaluated in microcebes. In both macaques and humans, microglial activation in white matter has been correlated with cognitive impairments (Gefen et al., 2019; Sloane et al., 1999). Unfortunately, the microcebes investigated in our study were not studied cognitively.

Unlike in white matter, HLA-DR+ microglia were sparsely detected in the parenchyma of cerebral cortex. This result was unexpected as previous studies using ferritin could identify microglia close to amyloid deposits in gray matter of microcebes (Trouche et al., 2023). In the only animal that exhibited amyloid deposits, HLA-DR+ microglia were not detected in the vicinity of amyloid plaques, however this could be due to the amyloid deposits being diffuse and sparsely distributed. Of note, this observation aligns with previous reports indicating that HLA-DR+ microglia preferentially associate with neuritic plaques in humans (Rogers et al., 1988) and are largely absent near diffuse plaques as also observed in macaques (Shah et al., 2010). Some parenchyma cortical regions adjacent to the corpus callosum, as the cingulate cortex, displayed HLA-DR+ microglia associated with myelin fibers extending from the corpus callosum as supported by the Luxol Fast Blue staining

### 4.2. Age-related increased HLA-DR+ perivascular macrophage

Here, we also investigated the presence of different macrophage populations: perivascular, meningeal and associated with the plexus choroid. Analysis based on cell counting displayed an age-related HLA-DR+ increase of perivascular macrophages in the cingulate, parietal and entorhinal cortices, as well as in the hippocampus, as previously reported in mice (Drieu et al., 2022; Mrdjen et al., 2018). Recent reviews highlighted the implication of perivascular macrophages in brain-associated pathologies such as infections or neurodegenerative diseases (Faraco et al., 2017; Kierdorf et al., 2019; Schonhoff et al., 2023). Their implication in Alzheimer’s disease progression has been suggested. Indeed, studies in rodent model of Alzheimer’s disease suggest that the perivascular macrophages are key cells to regulate the dynamics of the cerebral blood flow, contributing to brain clearance of small molecules such as Aβ (Drieu et al., 2022), and also limiting the development of tau pathology (Drieu et al., 2023).

Our study is the first one to describe increased number of HLA-DR+ perivascular macrophages in a primate. It is now acknowledged that, as microglia, they arise from the embryonic yolk sac (Wen et al., 2024). These cells have a wide range of functions including: phagocytosis, lymphatic drainage, maintenance of vascular integrity, and regulation of metabolic process (Wen et al., 2024). In physiological conditions, these cells are not replenished, while under pathological conditions, findings from preclinical models suggest that monocytes from the blood can contribute to the perivascular macrophage population (Wen et al., 2024). Further study will have to investigate if the age-related increase that we describe is related to a replenishment process occurring as part of aging. Some of the perivascular macrophages are HLA-DR-, but the HLA-DR+ ones are considered to be key for antigen presentation, playing a role in immune surveillance, exhibiting both phagocytic activity and antigen-presenting functions (Hawkes & McLaurin, 2009; Koizumi et al., 2019). Additionally, they contribute to the brain’s physiological response to circulating pro-inflammatory cytokines (Serrats et al., 2010). The aged-related changes in HLA-DR+ macrophages in the gray matter reported in this study could thus be related to activation of innate immune system during the aging process, although their exact function remains unclear.

### 4.3. Perspective for comparative pathology

The microcebe is a model used to study the impact of aging in primates (Languille et al., 2012), and several age-related changes reported in our study are comparable with changes reported in other primates. Indeed, consistently with our findings, age-related white matter microglial activation has been observed in other species as mice (Healy et al., 2024; Wong et al., 2005), rats (Perry et al., 1993), dogs (Tafti et al., 1996) or macaques (Sheffield & Berman, 1998; Sloane et al., 1999). Similarly, studies in humans reported increased microglial reaction in the white matter of older individuals (Amor et al., 2022; Gefen et al., 2019; Mattiace et al., 1990; McGeer et al., 1988; Rogers et al., 1988; Sobel & Ames, 1988). Few studies however outlined age-related diversity of white-matter microglia.

Regarding gray matter, as observed with HLA-DR labeling in the microcebe, few microglial cells were observed in cortical regions of humans during aging, in the absence of amyloid plaques (Mattiace et al., 1990; Rogers et al., 1988; Sobel & Ames, 1988). Similarly, lack of age-related cortical microglia activation has been reported in several non-human primates (marmosets (Rodriguez-Callejas et al., 2016), chimpanzee (Edler et al., 2017)). Results in macaques are more controversial with some studies suggesting cortical microglia activation (Peters et al., 1991; Robillard et al., 2016; Shobin et al., 2017)) while some other did not (Kanaan et al., 2010; Peters et al., 2008). Interestingly, microglial HLA-DR expression in gray matter was identified to be associated with injury, chronic disease states or pathological deposits (Mattiace et al., 1990; Rogers et al., 1988; Sobel & Ames, 1988).

## 5. Conclusion

This study characterizes for the first time microglia and their age-related changes in the whole brain of microcebes. Further research is needed to understand the origin of the two main changes reported here, *i.e.* white matter microgliosis in some old animals and increased perivascular macrophages at the level of cerebral cortex. The age-related changes reported here can be key factors leading to vulnerability to neurodegenerative diseases.

## 6. Acknowledgements

We thank the France-Alzheimer Association, ANR Primalz for funding this study. The MIRCen facility was funded by a grant from NeurATRIS: A Translational Research Infrastructure for Biotherapies in Neurosciences (“Investissements d’Avenir”, ANR-11-INBS-0011).

We thank the Agence Nationale de la Recherche (PrimAlz, ANR-22-CE14-0056), France-Alzheimer Association, Paris-Saclay University (Graduate School Life Sciences and Health) for funding this study. The MIRCen facility was funded by a grant from NeurATRIS: A Translational Research Infrastructure for Biotherapies in Neurosciences (“Investissements d’Avenir”, ANR-11-INBS-0011). LG was financed by the French Ministère de l’Enseignement Supérieure, de la Recherche et de l’Innovation.

## 8. Author contributions

L.D, L.G, M.D contributed to the study conception and design. F.P and S.L performed the microcebe euthanasia and collected brain samples. L.D, F.P, S.L performed immunohistological experiments. L.D conducted quantification. L.D and L.G analyzed the data. L.D, L.G, H.H., J.L.P., D.B, and M.D wrote the manuscript. M.D edited the manuscript. All authors commented on previous versions of the manuscript. All authors read and approved the final manuscript.

## 9. Data and code availability statements

The data that support the findings of this study are available from the corresponding author upon reasonable request.

