## Supplementary information for "Normal aging increases white matter microglial reaction and perivascular macrophages in the microcebe primate"

### Supplementary table 1

Microcebe characteristics. Both the middle-aged and old cohorts included male and female individuals. Medical information are reported when pertinent, otherwise nothing special to report (NSR) is written. FD: Found dead. SE: Euthanized for study purposes. AE: Euthanized to prevent suffering following an identified pathology.

| ID | Sex | Age (years) | Medical record | Death context |
|---|---|---|---|---|
| 1 | M | 2.3 | Sudden death - Undetermined cause of death despite autopsy | FD |
| 2 | M | 4.1 | NSR | SE |
| 3 | M | 4.8 | Tail infection | AE |
| 4 | F | 5.3 | Head shock | FD |
| 5 | F | 9.8 | Cataract | SE |
| 6 | M | 9.8 | NSR | SE |
| 7 | F | 9.8 | NSR | SE |
| 8 | M | 9,9 | NSR | SE |
| 9 | F | 10.3 | Cataract | SE |
| 10 | M | 10.3 | Eye ulcer | SE |
| 11 | F | 10.3 | NSR | SE |
| 12 | M | 10.3 | NSR | SE |
| 13 | M | 10.5 | NSR | SE |
| 14 | F | 10.5 | NSR | SE |
| 15 | M | 11 | NSR | SE |
| 16 | F | 11.4 | NSR | SE |
| 17 | F | 11.5 | Cataract | SE |

**Supplementary table 2**
Key resource table

| Reagent or Resource | Source | Identifier |
|---|---|---|
| **Antibodies** | | |
| Mouse anti-human HLA-DR Dilution: 1/250 | Dako | Cat#M0746 |
| **Chemicals and commercial assay or kit** | | |
| Impress Kit | Eurobio Scientific® | MP-7402 |
| ABC Vectastain® ABC-HRP kit | Vector Laboratories® | Cat#PK6100 |
| Luxol blue | FISHER SCIENTIFIC | 10348250 |
| Carbonate de lithium | VWR | MOLE13183442-100G |
| Bovine serum albumin (BSA) | Sigma-Aldrich® | Cat#A7906 |
| Citrate Buffer 10X, pH 6.0 | Diagnostic BioSystems® | Cat#924 |
| Cresyl violet | Merck | Cat#10510-54-0 |
| DAB Substrate kit, Peroxidase (HRP), with Nickel | Vector Laboratories® | Cat#SK4100 |
| Dulbecco's phosphate saline (DPBS) 1X | Gibco™, ThermoFisher | Cat#14190094 |
| Ethanol absolute | VWR | Cat#83813360 |
| Ethylene glycol | Carlo Erba | Cat#346502 |
| Eukitt® mounting medium | Sigma-Aldrich® | Cat#03989 |
| Foetal bovine serum | Sigma-Aldrich® | Cat#F7524 |
| Formic acid | VWR® | Cat#BDH4554 |
| Glycerol | Fisher | Cat#12144481 |
| Horse serum | Gibco™, ThermoFisher | Cat#16050122 |
| Hydrogen peroxide 30% | Sigma-Aldrich® | Cat#H1009 |
| Ketamine | Imalgène® 1000, Merial | |
| Lidocaine | 0.5% Xylovet®, Ceva Santé Animale | |
| Liquid nitrogen | Air products, France | |
| Normal goat serum (NGS) | Sigma | Cat#G6767 |
| Paraformaldehyde, PFA | Sigma | Cat#P7148 |
| Pentobarbital | Exagon®, Axience | |
| Phosphate buffer solution, 1 M , pH 7.4 | Sigma-Aldrich® | Cat#P3619 |
| Phosphate Buffered Saline (PBS), pH 7.4 | Sigma-Aldrich® | Cat#806552 |
| Sodium chloride (NaCl) | Sigma-Aldrich® | Cat#S9888 |
| Sucrose | Sigma-Aldrich® | Cat#S0389 |
| Tris-HCl | Merck | Cat#252859-500G |
| Triton X-100 | Sigma-Aldrich® | Cat#X100 |
| Xylazine | 2% Rompun®, Bayer Healthcare | |
| Xylene | VWR Chemicals | Cat#28973363 |
| **Equipements** | | |
| 60x water-immersion objective | Nikon, melville, ny, usa | |

| | | |
|---|---|---|
| 40x water-immersion objective | Leica | |
| Argon laser | Nikon, melville, ny, usa | |
| Axio Scan.Z1 | Zeiss® | |
| Centrifuge | Multifuge X1R Heraeus Thermoscientific | |
| Confocal optical microscope (TCS SPE) | Leica DMI6000 | |
| EPC 10 Amplifier Patchmaster Multi-channel | HEKA Elektronik Dr. Schulze gmbh, Wiesenstrasse, Germany | |
| Heating box | Phymep | |
| Heating pad | Phymep | |
| Microtome | Leica Vt1200 blade | |
| OptimaTM TL 100 Ultracentrifuge | Beckman | |
| Perfusion pump | Fisher Scientific | Cat#1170-5369 |
| SM2400 microtome | Leica Microsystems | |
| Spark plate reader | Tecan | |
| SW32ti swinging bucket rotor | Beckman | |
| Ti C2 confocal microscope | Nikon, melville, ny, usa | |
| TLA-100.2 fixed angle rotor | Beckman | |
| DMI6000 confocal optical microscope | TCS SPE, Leica | |
| **Experimental models: Organisms / Strains** | | |
| Microcebus Murinus (WT) | Brunoy - MNHN | |
| **Material** | | |
| 6 well plates | | |
| 24-well plates | Falcon; Beckton Dickinson | |
| Superfrost Plus slides | Thermo-Scientific® | |
| **Software and algorithms** | | |
| ImageJ Software | https://imagej.nih.gov/ij/download.html | |
| QuPath v0.4.3 software | https://qupath.github.io/ | |
| R Studio 4.3.3 | https://www.r-project.org/ | |
| R-package Rcmdr | https://cran.r-project.org/web/packages/Rcmdr/index.html | |
| GraphPad Prism software 9 | https://www.graphpad.com/ | |
